# Genetic diversity and population structure among goat genotypes in Kenya

**DOI:** 10.1101/2020.07.06.189290

**Authors:** Ruth W. Waineina, Kiplangat Ngeno, Tobias O. Otieno, Evans D. Ilatsia

## Abstract

Population structure and relationship information among goats is critical for genetic improvement, utilization and conservation. This study explored population structure and level of gene intermixing among four goat genotypes in Kenya: Alpine (n = 30), Toggenburg (n = 28), Saanen (n = 24) and Galla (n = 12). The population structuring and relatedness were estimated using principal component analysis utilizing allele frequencies of the SNP markers. Genotype relationships were evaluated based on the calculated Reynolds genetic distances. A phylogenetic tree was constructed to represent genotype clustering using iTOL software. Population structure was investigated using model-based clustering (ADMIXTURE) Genotypes relationships revealed four distinctive clusters: Alpine, Galla, Saanen and Toggenburg. The ADMIXTURE results revealed some level of gene intermixing among Alpine, Toggenburg and Saanen with Galla. Saanen goats were the most admixed genotype with 84%, 7% and 4% of its genome derived from Galla, Alpine and Toggenburg respectively. Alpine and Toggenburg goats shared some associations with the Galla goat; 10% and 1% respectively. The association of Galla with other genotypes was anticipated since Galla goat was used as the founder population for crossbreeding with Saanen, Alpine and Toggenburg breed. The genetic variations among the goat genotypes observed, will provide a good opportunity for sustainable utilization, conservation and future genetic resource improvement programs in goat genotypes in Kenya.

## Introduction

Goats are known to be the most adaptable and widespread across various geographical conditions, ranging from the mountains to the deserts and tropics of Africa. The roles of goats in supporting rural household economies in developing countries are well documented [1-3]. They form important sources of food and nutritional security through the supply of milk and meat, income generation through sale of surplus stock and insurance against unforeseen risks and other non-tangible cultural values [4-6]. Recent studies have shown that goats farming is one of the alternative climate-smart agricultural practices that could build farmers resilience to climate change-related challenges [7]. The diminishing land sizes in the medium to high potential areas due to human population pressure, expansion of urban areas and climate change-related challenges are triggers to alternative farming practices such as intensive dairy goat production [8]. This is because dairy goat production offers more multi-functionality, flexibility and adaptability to varied production conditions.

In Kenya, dairy goat production has mainly been supported by exotic breeds (Toggenburg, Anglo-Nubian, German Alpines, Saanen and Boer) and their crosses with selected local breeds (Galla and small East African goat) [9-11]. The exotic breeds were introduced to various parts of the country by the Government of Kenya (GoK) and Non-Governmental Organizations (NGOs). The ultimate aim has been to increase their productivity through appropriate husbandry and disease interventions [12] and targeted breeding strategies such as crossbreeding [12, 13].

Crossbreeding has been the breeding strategy of choice to improve the productivity of goats under various production systems [12, 14]. This has resulted in an increase in population sizes of crossbred goats especially in the target projects areas and other areas apart from the original entry points in the country [12, 15]. However, the increase in population sizes did not necessarily correspond to enhanced productivity but rather reflected the many numbers of households who were striving to support their livelihoods through goat farming [15-17].

There was limited technical capacity on the farmers’ side on how to manage the rather complex crossbreeding programs, a fact that could have had a bearing on the sustainability of such initiatives in the long term [15-17]. The net result of this has been the un-systematic crossing of the existing poulation, poor flock management, lack of records to support decision making and generally lack of simplified breeding programs to guide in genetic improvement of goats in the country [18]. Currently, admixtures of exotic and local goats are reared as dairy goats in different parts of the country under different production systems. There is a huge source of genetic diversity in the current goat populations in Kenya. This is as a result of unsystematic crossbreeding and lack of records keeping by most of the smallholder farmers. This calls for the need to characterize, conserve and utilize sustainably under various production systems. It is important to determine genetic diversity in populations because it provides the basis for natural and artificial selection [19].

To measure and describe genetic diversity in the animal genetic resource, phenotypic and molecular characterization tools are used as a starting point to understand the animal resources and make use of them sustainably [20]. This includes all information on breed origin, development, structure, population, quantitative and qualitative characteristics in defined management and climatic conditions [21, 22].

Molecular characterization using genetic markers are powerful tools which can be realised in many applications in a breeding program. For instance, it can be used to characterize the genetic variability within and genetic distance between breed populations, genomic selection, parentage verification and genetic diversity preservation [23, 24]. This can be done by using categories of genetic markers to detect polymorphism in nuclear DNA.

Microsatellite and single-nucleotide polymorphisms [SNPs] are the most commonly used markers in animal breeding related fields [25]. Microsatellite, however, has limitations in genetic diversity which includes; null alleles [24], homoplasy [26, 27] and linkage disequilibrium [28]. SNP several advantages relative to Microsatellite, among others, SNP is highly reproducible and very informative [29] and markers can represent either neutral or functional genetic diversity [30]. GoatSNP50 Bead Chip (Illumina, Inc. San Diego, CA 92122 USA) which was developed from SNP loci detected by whole-genome sequencing of six different goat breeds according to Tosser-Kloppet et al [31] is available. This has made SNPs markers to be the most popular and advanced technology in molecular breed characterization in goats. Additionally, its robustness, low genotyping costs, automatic allele calling and capability to interrogate the goat genome at high resolution [32] demonstrate practicability to implement genomic characterization in goats.

In Kenya, there has been no deliberate effort to understand the genetic diversity and population admixture among the goat genotype populations by use of SNPs makers. This study, therefore, investigated genetic diversity, population structure and admixture among goat genotypes in Kenya. The results from this study will provide information which can be used in facilitating management efforts in conserving and utilizing the various goat genetic resources sustainably.

## Materials and Method

### Study area

The study was conducted from August to September 2018 in three Counties in Kenya as presented in Fig 1. They include, Nyeri (Mukurweini Sub-County), Meru (Central Imenti Sub-County) and Homa Bay (Homa Bay town Sub County) located in the Central, Eastern and Western regions of Kenya, respectively. These areas were selected because they were the entry points of different exotic dairy goat breeds in Kenya. Mukurweini sub-county lies in the Upper midlands (UM2), also known as the main coffee zone, is 1460-1710 meters above sea level and receives 950-1500mm of mean annual rainfall. Central Imenti is in the upper highlands (UH) at an altitude ranging between 1830-2210 meters above sea level and has an average precipitation of 800-2600mm annually. Homa Bay town sub-county lies in the lower midlands (LM2).

**Fig 1.**
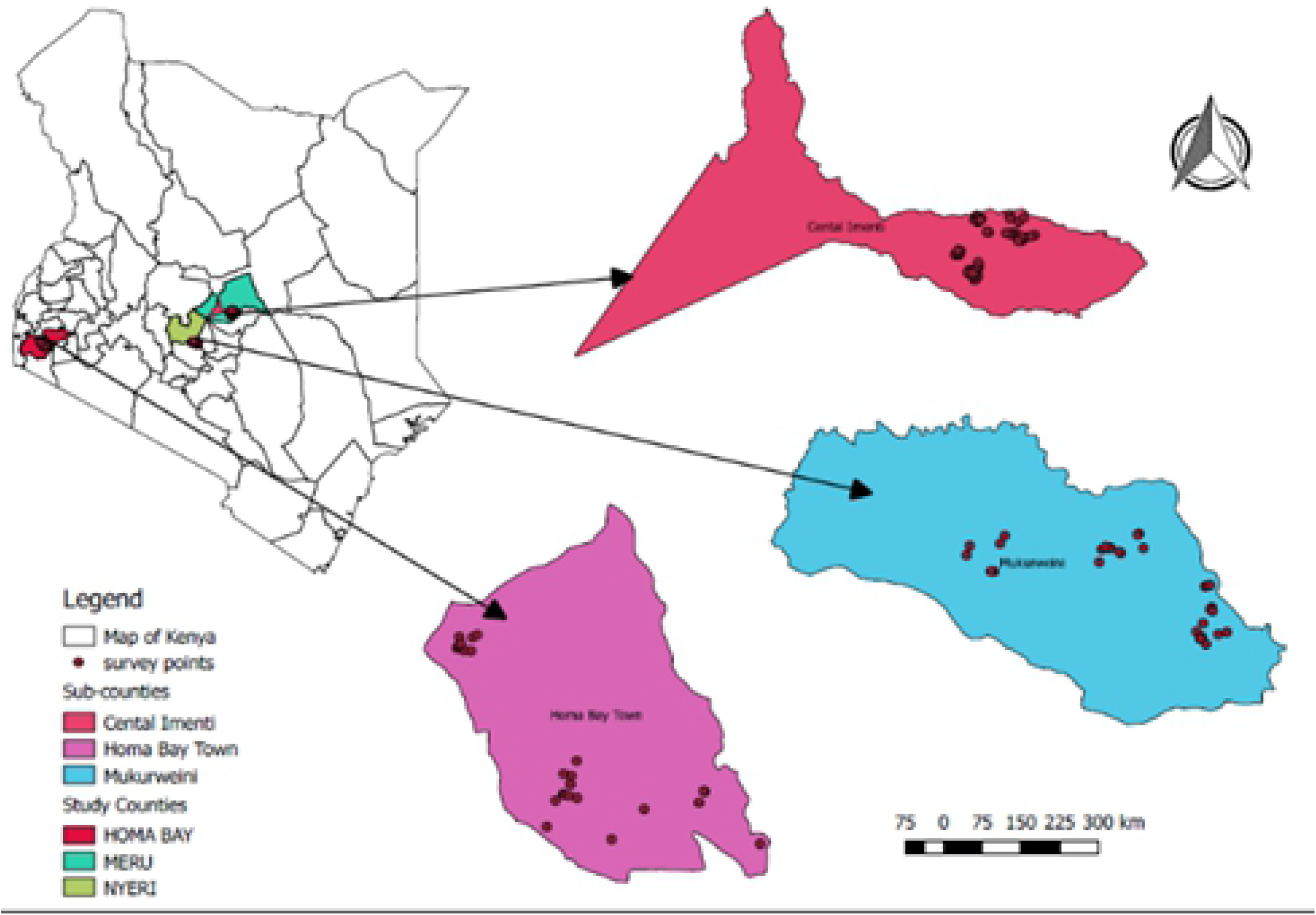
Map of Kenya showing area sampled within the selected sub-counties.

### Animal resources and sampling

Goat keeping households were purposively selected based on the following criteria: being a member of dairy goat farmer group and owns at least a mature doe which matches the genotype kept by the farmer group in the said County of study. To ensure the representativeness of sampling, for each breed, unrelated animals were selected from various farms across the designated Counties. The Galla goat breed, however, did not follow the criteria because they were from the Government breeding station where record keeping was used to avoid sampling related animals.

A total of 96 animals including three exotic genotypes (Toggenburg = 30) (Alpine = 28), (Saanen = 24) and one indigenous (Galla = 12) goats were incorporate in this study. The Toggenburg and Alpine were found in Eastern and Central Kenya under Meru Goat Breeders Association (MGBA) and Dairy Goat Association of Kenya (DGAK), respectively. Saanen goats were found in Homa-Bay. Galla goats were sampled from sheep and goat Government station in Naivasha.

Whole blood (10ml) was collected from the jugular vein into Vacutainer tubes with Ethylenediaminetetraacetic acid (EDTA) as an anticoagulant. The blood was stored at -20°C for two months before genomic DNA extraction. The procedure of blood sampling followed [33] guidelines. To allow for possible loses, mistyping and missing values, two adult does per household was sampled. From each animal, a duplicate was collected and kept separately during transportation and storage. For each sample, the following information was collected: sex of the animal, basic pedigree information, size of the flock, breed, any relevant phenotypic feature and a photograph of the goat.

The study was conducted in strict accordance with the recommendations for the Institute of Primate Research (IPR) ethical guidelines on animal care and use of laboratory animals and also for the collaborating organizations. The protocol was approved by the committee on the ethics of Animal Experiments of the University of Egerton in Kenya. A qualified veterinary officer collected the whole blood following FAO (2011) guidelines to reduce pain and discomfort to a minimum.

### DNA extraction and genotyping

Genomic DNA was extracted from the whole blood using the DNeasy Blood and tissue kit (Qiagen®, Hilden, Germany). Genomic DNA was quantified using Nanodrop Spectrophotometer (Nanodrop ND-1000) and genotyped using the GoatSNP50 Bead Chip (Illumina, Inc. San Diego, CA 92122 USA) developed by the International Goat Genome Consortium (IGGC) [31].

### Data analysis

The following filtering was applied to raw reads; SNPs with less than 95% call rate, less than 0.05 Minor Allele Frequency (MAF), Hardy Weinberg Equilibrium (HWE) P<0.001 and more than 10% missing genotypes were filtered using Plink v1.9 [34]. Genetic diversity basic indices which include the proportion of polymorphic markers (P_N_) inbreeding coefficient, observed and expected heterozygosity were calculated within genotypes using PLINK [34]. The population structuring and relatedness were estimated by principal components analysis (PCA) from the R package SNPRELATE [35]. Besides, the population structure was investigated using model-based clustering ADMIXTURE 1.3.0 software [36]. This helped to deduce the true number of genetic population (cluster or K) between the four goat genotypes. ADMIXTURE cross-validation (CV) procedure determined the optimal cluster or K-value. The K-value with the lowest CV error was selected as the optimal value. Phylogenetic tree based on Reynolds genetic distances representing genotypes relationships among ‘goat genotypes was visualized using iTOL software [37].

## Results

Quality control procedure on the 53,347 SNPs included in the SNP chip excluded a total of 3,542 SNPs retaining 9,805 SNPs for downstream analysis (Table 1). Out of the SNPs excluded, 2235 as a result of less than 0.1 missing per SNP. About 663 SNPs significantly deviated from HWE (P < 0.001) and 644 had MAFs less than 0.05. Galla genotype showed the highest number of SNPs excluded in total (0,688), whereas Alpine revealed the lowest number of SNPs excluded (5,129).

**Table 1.** Goat genotype, number of goats, number of exclude and remaining SNPs and number of goats.

| Genotype | N | Excluded SNPs |  |  |  |  | Remaining SNPs | Remaining samples |
| --- | --- | --- | --- | --- | --- | --- | --- | --- |
|  |  | MIND | GENO | HWE | MAF | Total |  |  |
| Saanen | 24 | 0 | 2453 | 38 | 3658 | 6149 | 47198 | 24 |
| Alpine | 29 | 1 | 2496 | 47 | 2586 | 5129 | 48218 | 28 |
| Galla | 12 | 0 | 2413 | 26 | 8249 | 10688 | 42659 | 12 |
| Toggenburg | 31 | 1 | 2345 | 50 | 4690 | 7085 | 46262 | 30 |
| Merged | 96 | 2 | 2235 | 663 | 644 | 3542 | 49805 | 94 |

N, number of animals; MIND, genotype missing ( $< 0.1$ ), GENO, SNP missing ( $< 0.15$ ), MAF, minor allele frequency; HWE, Hardy-Weinberg equilibrium ( $P$ -value $< 0.001$ ).

### Genetic diversity

The four indices (polymorphic SNPs, mean allele frequency, observed and expected heterozygosity and inbreeding coefficient) of genetic diversity were calculated within each genotype (Table 2). The assessment of the proportion of SNPs that exhibited both alleles within each genotype indicated high levels of diversity. The percentage of within-genotype polymorphic SNPs ranged from 94% to 80.7%. The highest values of polymorphic were observed in Alpine (94.6%) and Saanen (92.2%) and both were involved in SNP chip discovery panel. The lowest proportion of polymorphic loci (P_N_) was detected in Galla genotype (80.7%). Across all the loci, the lowest MAFs was found in Galla (0.291) and highest in Alpine (0.323).

**Table 2.**
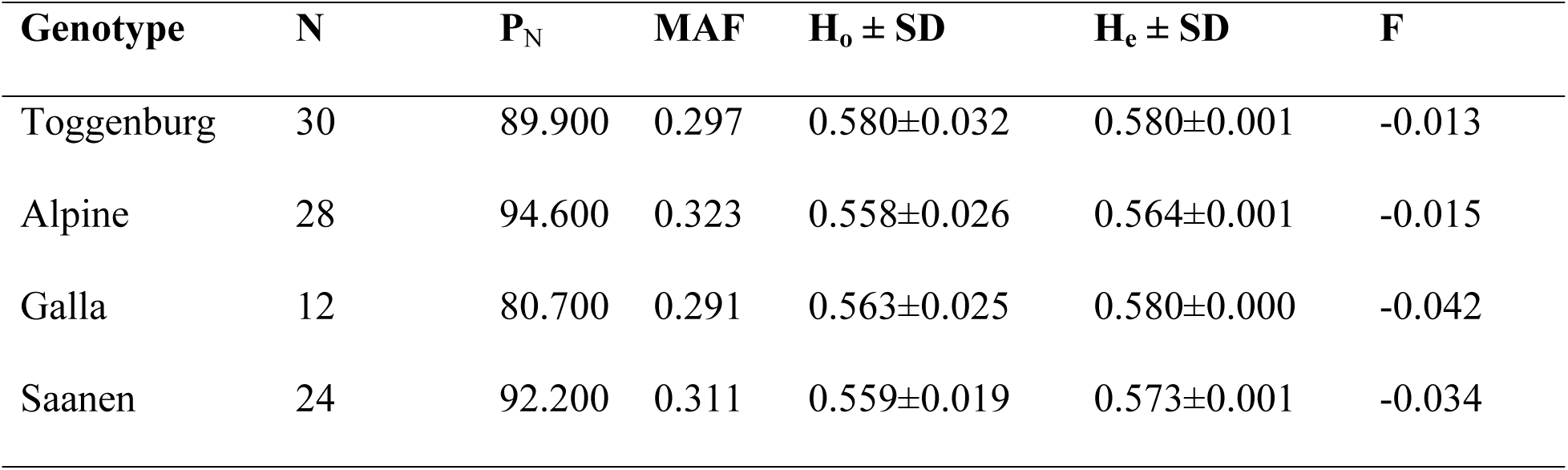
Proportion of polymorphic SNPs (P_*N*_) mean allele frequency (MAF), Observed (*H*_*o*_) and expected (*H*_*e*_) heterozygosity and inbreeding coefficient (F)for the goat genotypes.

| Genotype | N | $P_N$ | MAF | $H_o \pm SD$ | $H_e \pm SD$ | F |
| --- | --- | --- | --- | --- | --- | --- |
| Toggenburg | 30 | 89.900 | 0.297 | 0.580±0.032 | 0.580±0.001 | -0.013 |
| Alpine | 28 | 94.600 | 0.323 | 0.558±0.026 | 0.564±0.001 | -0.015 |
| Galla | 12 | 80.700 | 0.291 | 0.563±0.025 | 0.580±0.000 | -0.042 |
| Saanen | 24 | 92.200 | 0.311 | 0.559±0.019 | 0.573±0.001 | -0.034 |

Results revealed differences in genetic diversity between genotypes. The expected heterozygosity was relatively greater than observed heterozygosity (*H*_*e*_ > *H*_*o*_). In comparison, Alpine recorded the lowest observed heterozygosity (H_o_ = 0.558 ± 0.026) while Toggenburg had the highest (H_o_ = 0.580 ± 0.032). Inbreeding coefficients for all the genotypes were negative and ranged between - 0.013 (Toggenburg) and -0.042 (Galla).

### Population structure analysis

Principal components analysis was used to cluster goats and explore the association among individuals and genotype groups. In Fig 2, the principal component, Eigenvector 1 (EV1) which accounts for 15.2% of the total variance separated the various goat genotypes into differentiated clusters. The second principal component (EV2) accounts for 14.1% of the total variance, splitted the goat genotypes into four clusters (Alpine, Saanen, Galla and Toggenburg clusters). One outlier, was, however, observed for the Saanen population mixing with Alpine population.

**Fig 2.**
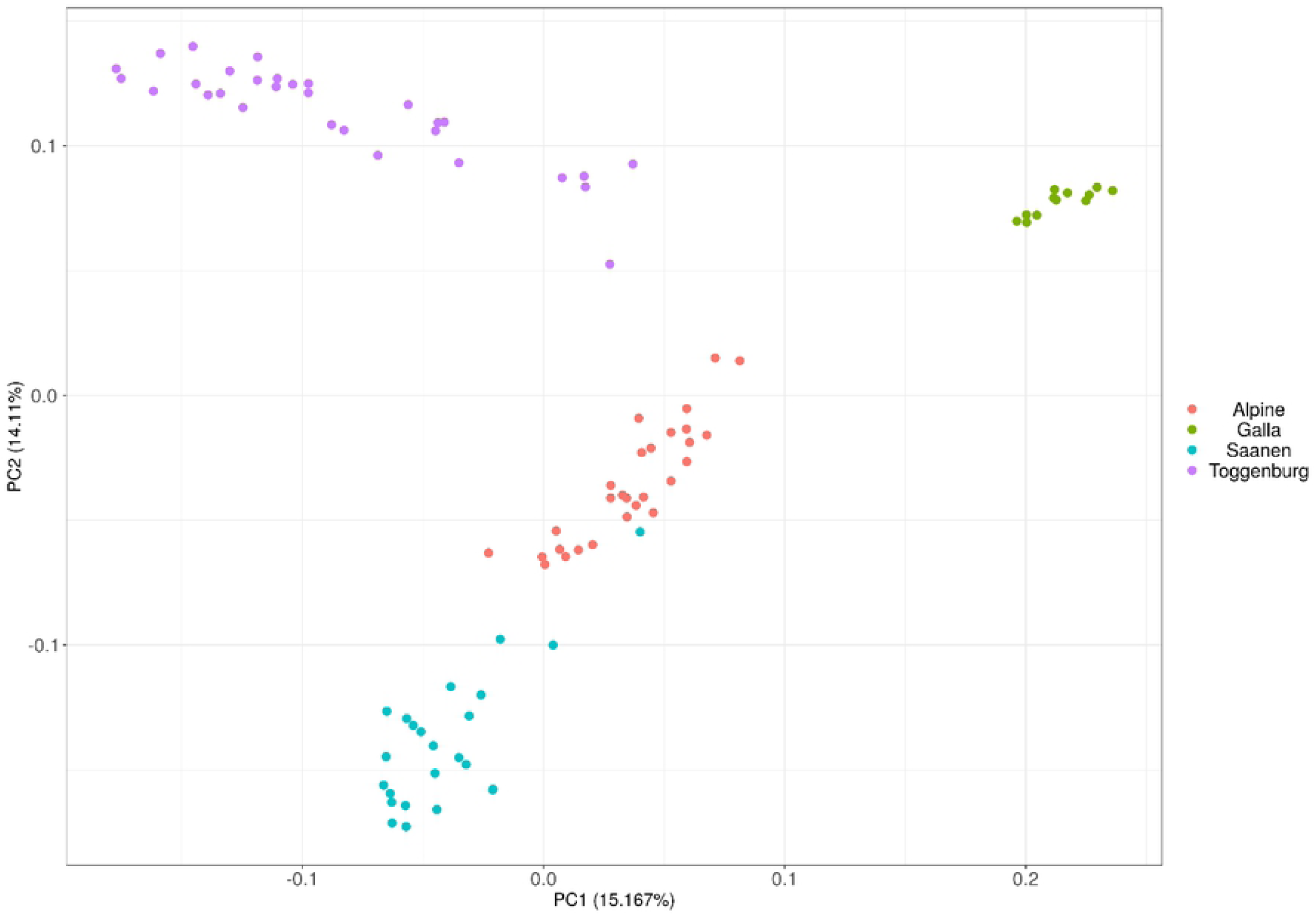
Principal components analyses plot based on SNP array data of goat genotypes.

To examine admixture between the genotypes, model-based clustering was performed and the true number of genetic population (cluster or K) between the four goat genotypes were deduced using ADMIXTURE cross-validation (CV) procedure. The K-value with the lowest CV error was K = 4 and was selected as an optimal number of ancestral populations (Fig 3).

**Fig 3.**
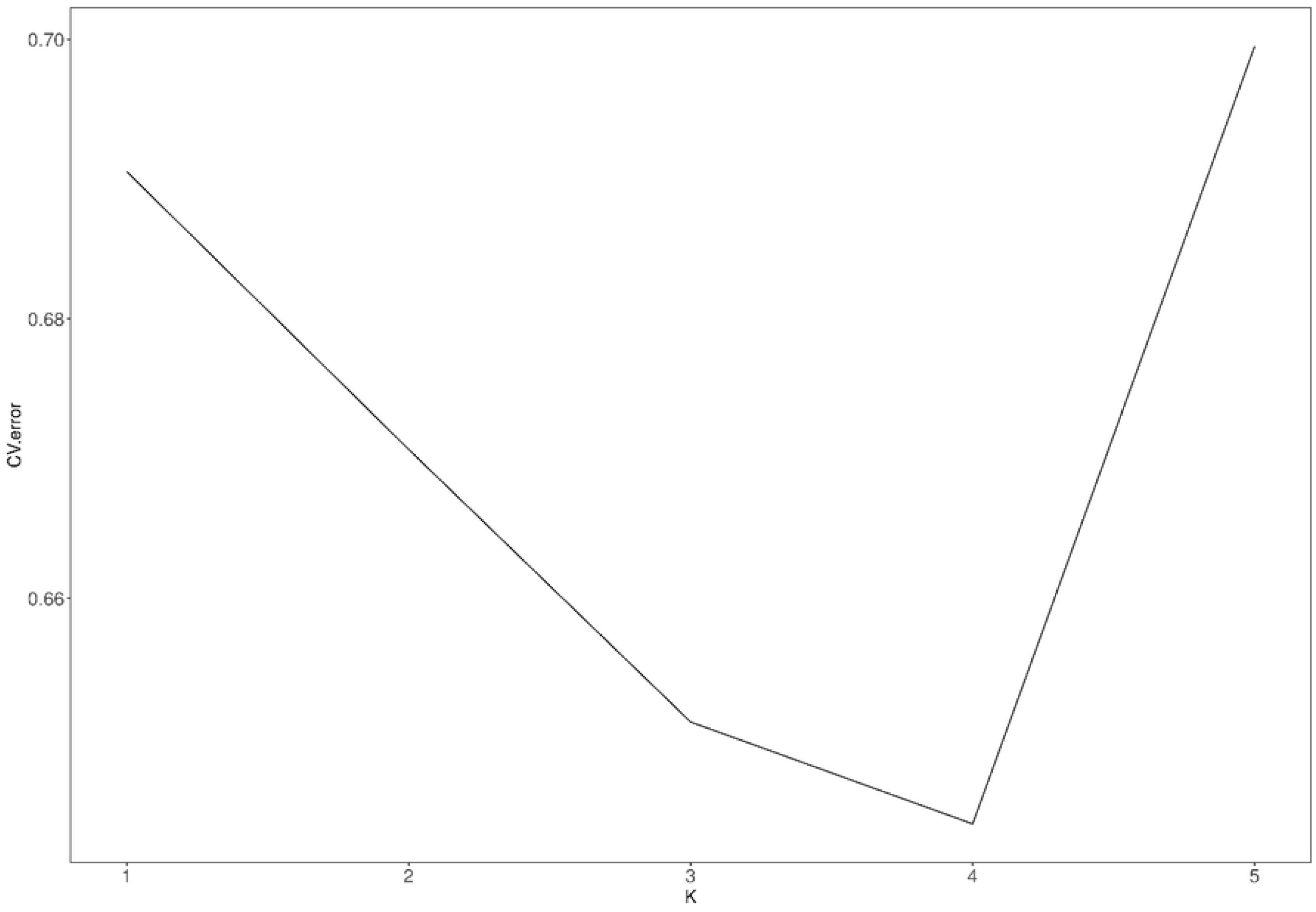
A cross-validation plot indicating the choice of optimal K-value.

Population structure plot (Fig 4) showed proportions of ancestral populations for every genotype (Alpine, Galla, Saanen and Toggenburg) for K = 2 to K = 4. At K = 2, Galla goats were separated from the other three goat genotypes (Toggenburg, Saanen and Alpine) Moreover, Galla goats largely do not carry ancestral components present in Saanen, Alpine and Toggenburg goats (shown in light blue, Fig 4). At K = 3, Alpine and Saanen goats carry ancestral components largely absent from either the Galla or Toggenburg goats. This most likely reveals the past genetic contribution of a breed that has not been included in this study. At K = 4, it revealed that Galla had the least level of admixture, whereas, Toggenburg, Alpine and Saanen demonstrated some signs of admixture with Galla.

**Fig 4.**
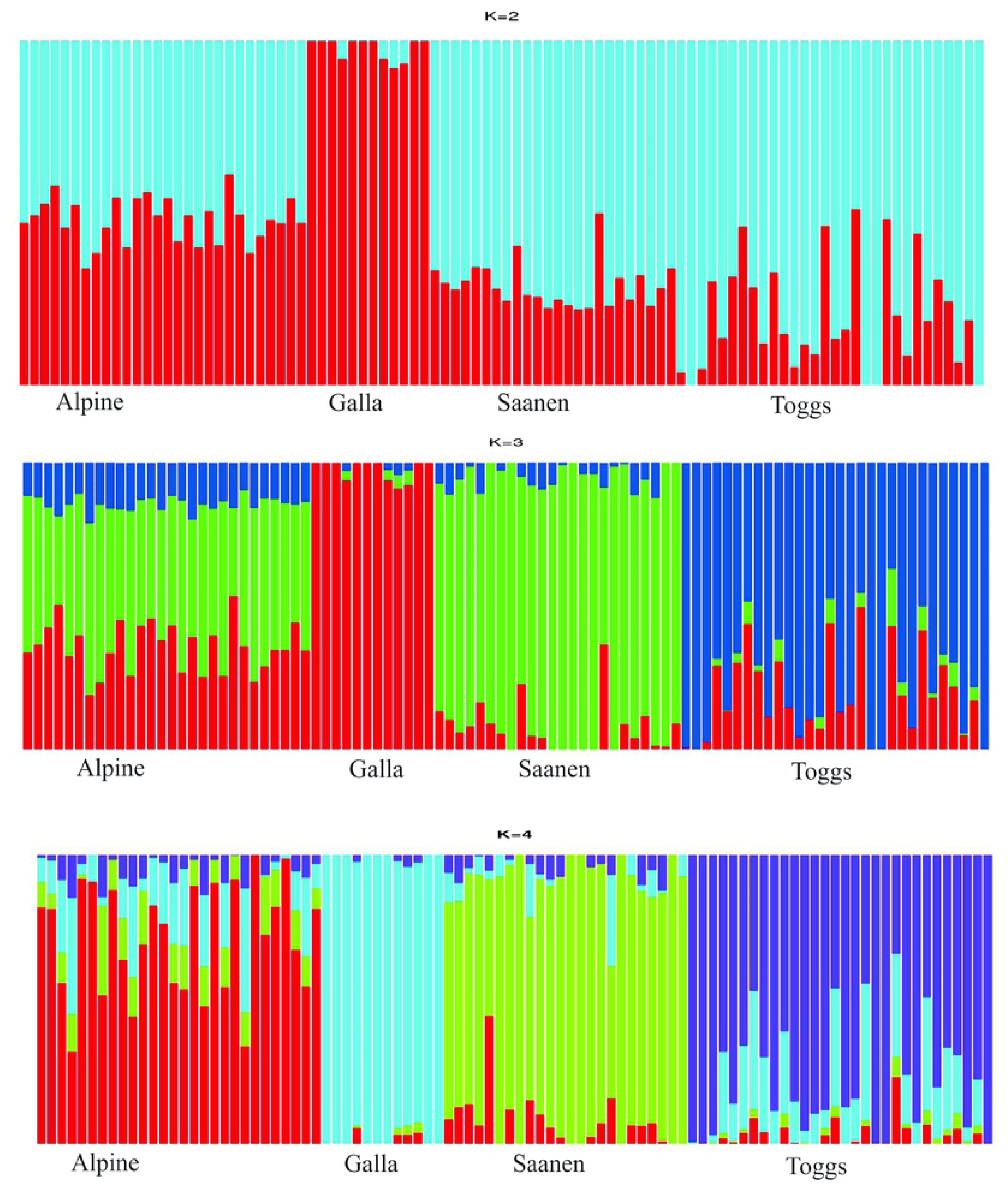
Population structure plots showing proportions of ancestral populations for every individual (Alpine, Galla, Saanen and Toggenburg (Toggs) for K = 2 to K = 4].

The proportions of individuals in each of the genotype in the four most likely clusters estimated by the ADMIXTURE as shown in Table 3 correspond to the four genotypes included in Figs 2 and 3. Results revealed that 71% of Alpine genotype were allocated to cluster one, one percent (1%) of Galla were assigned to cluster two with 97% of its genome assigned to cluster three, five percent (5%) of Saanen were in cluster three with 84% of its genome assigned to cluster two. On the other hand, 78% of Toggenburg was assigned to cluster four with three percent (3%) of its genome allocated to cluster one.

**Table 3.**
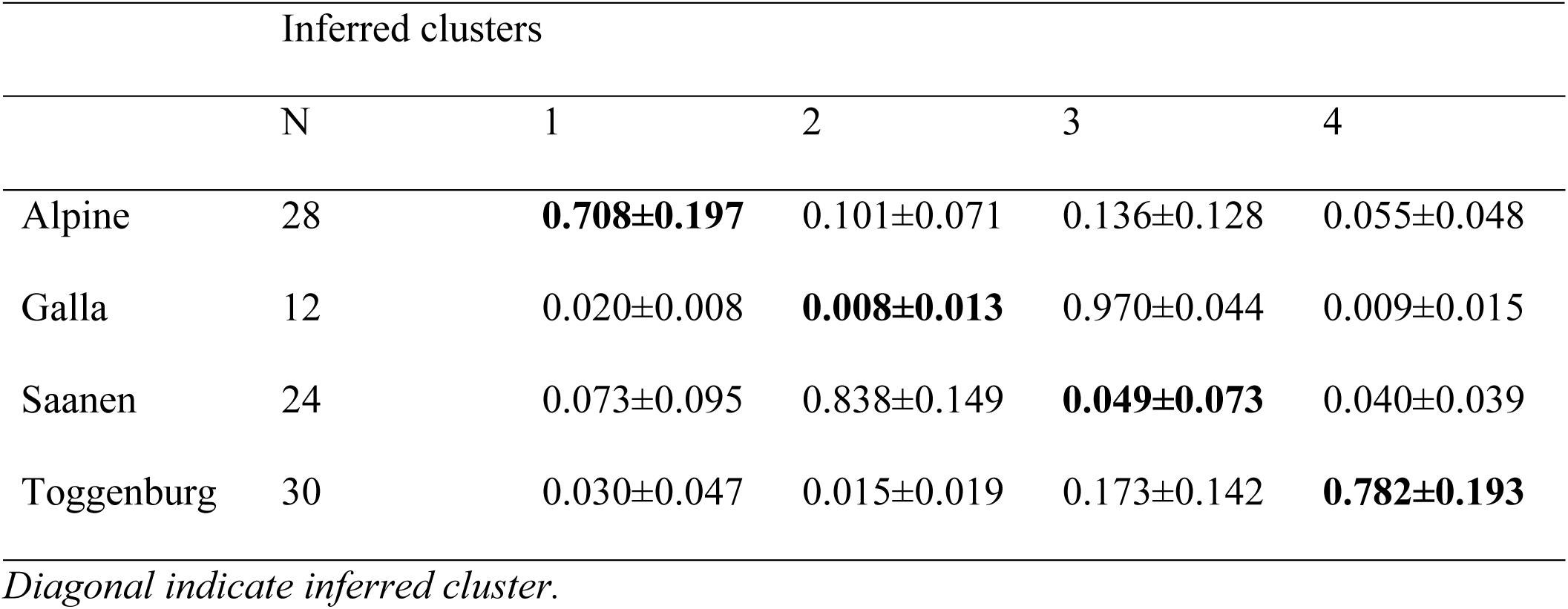
Proportion of individuals of goat genotypes in each of the four clusters estimated by ADMIXTURE.

|  | N | 1 | 2 | 3 | 4 |
| --- | --- | --- | --- | --- | --- |
| Alpine | 28 | **0.708±0.197** | 0.101±0.071 | 0.136±0.128 | 0.055±0.048 |
| Galla | 12 | 0.020±0.008 | **0.008±0.013** | 0.970±0.044 | 0.009±0.015 |
| Saanen | 24 | 0.073±0.095 | 0.838±0.149 | **0.049±0.073** | 0.040±0.039 |
| Toggenburg | 30 | 0.030±0.047 | 0.015±0.019 | 0.173±0.142 | **0.782±0.193** |

*Diagonal indicate inferred cluster.*

Genotype relationships were evaluated by computing the genetic distance between all pairwise combinations of individuals (D) from the average proportion of allele shared. Based on the calculated Reynolds genetic distances, a phylogenetic tree was constructed to represent genotype clustering (Fig 5). Population fitting to the same genotype clustered together as inferred by the identity by state (IBS) distance. The results revealed four distinctive clusters: Alpine, Galla, Saanen and Toggenburg (Toggs). This also confirms the results of the principal component and ADMIXTURE analysis.

**Fig 5.**
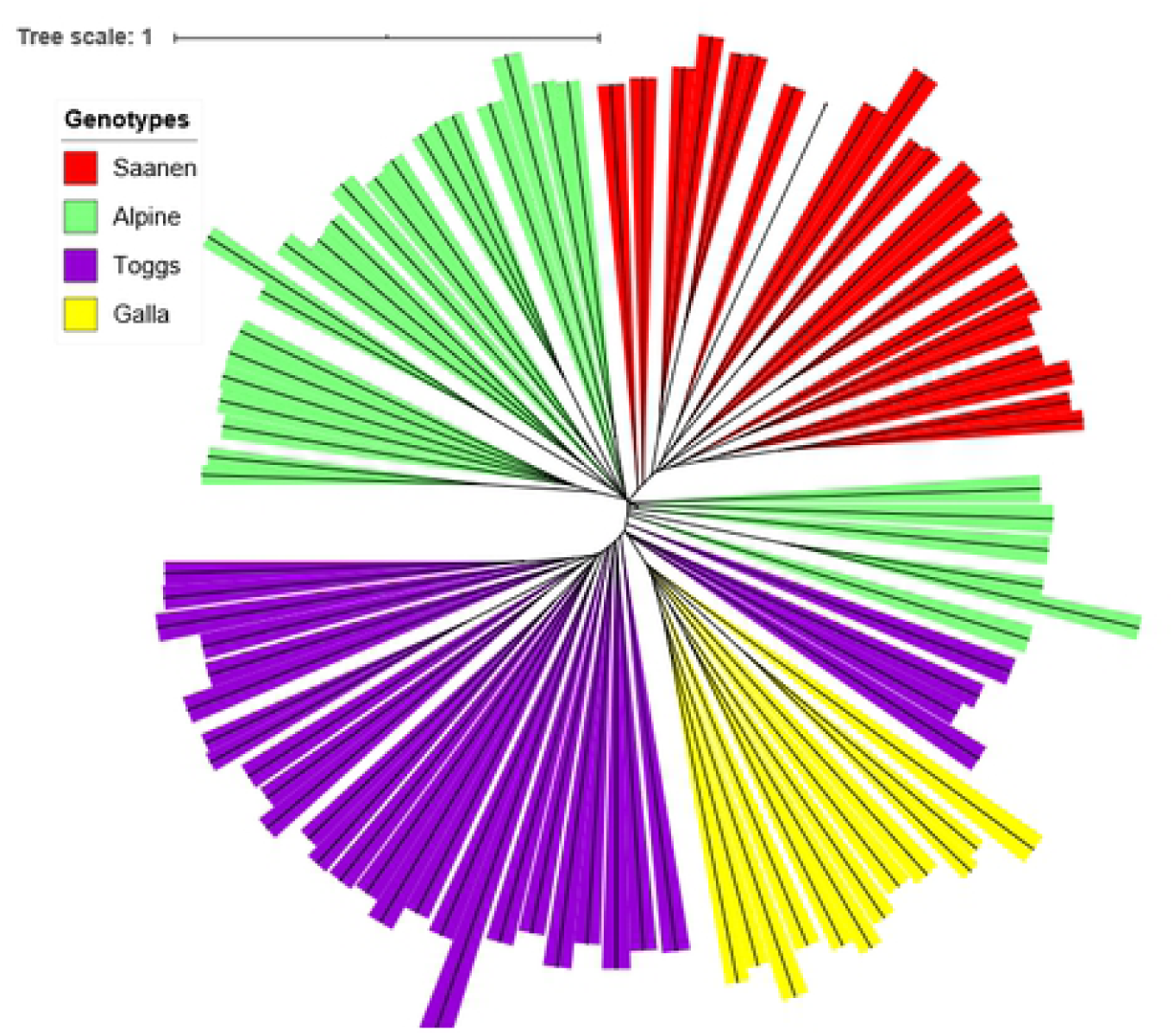
Phylogenetic tree based on Reynolds genetic distances representing genotypes relationships among goat populations.

## Discussion

### Genetic diversity

Livestock has been exposed to various forces that contributed to the genetic diversity underlying phenotypic dissimilarities ever since domestication. These forces include; natural selection, artificial selection for specific traits, migration, genetic drift and inbreeding [23, 38]. In Kenya, Toggenburg, Saanen and Alpine were imported to crossbreed with the local goats [Galla and small East African] to improve the milk productivity of local goats under various production systems. They were kept in different geographical locations as isolated populations and subjected to separate breeding objectives for some decades. With the availability of genomic technology, the next step was to investigate genetic diversity and relationships of the different populations/genotypes.

Genotyping with GoatSNP50 Bead Chip revealed some levels of diversity within the goat genotypes in this study. In each genotype, less than 80% of SNPs exhibited polymorphism, and gene diversity ranged from 0.558 to 0.580 (Table 2). A large number of polymorphic SNPs were detected for Alpine and Saanen genotypes; this was expected because sequenced data from Alpine and Saanen were included in the 50K SNPs panel discovery [31]. Other results using the different number of samples and goat breeds showed > 93% of polymorphism [39, 40]. It is, however, very difficult to compare and conclude on the estimates of SNPs stated as polymorphic by other authors because the number of samples genotyped per breed and proportion of genotyped samples used for SNP discovery varied.

The diversity amongst the four genotypes showed Galla had the lowest polymorphism in comparison with the other genotypes. The polymorphic variance is dependent on the history of each genotype. Apart from Galla goats, the other three genotypes were as a result of crossbreeding with local goats (Galla and small East African). Therefore, each genotype may contain genetic contributions from various breeds, thus revealing high polymorphism than Galla goats. In contrast, Galla goats were sampled from the Government breeding station with reputable pedigree records but still maintained levels of diversity (P_N_ = 81%). This means that genetic evolution can be expected to continue and no fear of inbreeding if selection pressure for target traits is maintained. Additionally, genetic variation to a population allows some individuals to adapt to the environment whereas the continuous existence of individuals is maintained. Inbreeding coefficient observes the probability that the alleles have come from a common ancestor. The negative inbreeding coefficient values in the four genotypes indicated limited inbreeding. It could also mean in this study that many heterozygotes were observed although the sample size for the four genotypes was small.

The observed heterozygosity was lower than the expected heterozygosity (*H*_*o*_ < *H*_*e*_) in the three genotypes apart from Toggenburg which recorded the same value for both observed and expected heterozygosity. The difference between the observed and expected heterozygosity was small which may not be due to inbreeding but a Wahlund effect. The observed heterozygosity in the current study is from a sample of individuals gotten from a structured population even though all subdivisions are in Hardy-Weinberg equilibrium. Over and above the semi and intensive production systems practised by smallholders, there is the presence of artificial selection, gene flow and non-random mating hence not holding the law of HWE in these populations. In this study the observed and expected heterozygosity for Alpine (*H*_*o*_ = 0.558; *H*_*e*_ = 0.564), Saanen (*H*_*o*_ = 0.559; *H*_*e*_ = 0.573) and Toggenburg (*H*_*o*_ = 0.580; *H*_*e*_ = 0.580) were higher than those stated in Canada for Alpine (*H*_*o*_ = 0.385; *H*_*e*_ = 0.388), Saanen (*H*_*o*_ = 0.379; *H*e = 0.382) and Toggenburg (*H*_*o*_ = 353; *H*_*e*_ = 336) [41]. Moreover, Saanen in Italy recorded the same trend as in Canada (*H*_*o*_ = 0.41; *H*_*e*_ = 0.41) [42]. Differences in effective population sizes, length of isolation, selection and breeding management practices in the various production system, may be the cause of variances.

Toggenburg and Galla genotypes, had the highest expected heterozygosity. This could be explained by the observed type of crossbreeding program practised by farmers keeping Toggenburg genotypes resulting into an admixed population. On the hand, organized breeding strategies by use of artificial selection is practised for Galla goat under the Government breeding station. This has resulted in genetic variability and lack of inbreeding for the populations.

### Population structure and relationship

Principal component, admixture and phylogenetic tree analysis of population structure confirmed distinctiveness among the goat genotypes (Saanen, Galla, Toggenburg and Alpine) in Kenya. This can be explained by the demographic history of these genotypes that have been reared for a long time in a separate geographic locations. Although goats from each genotype clustered separately, model-based clustering revealed that admixture has occurred and genetic links exist between the genotypes due to crossbreeding program

Saanen goat shared high genetic associations with Galla goat, with 84% of its genes resulting from Galla goat, although the PCA differentiated the two genotypes into well-separated clusters. The differentiation might be as a result of Saanen genotype having genetic links with other genotypes but, Galla goat is not admixed. Saanen was the most intermixed genotypes in this study with 84%, 7% and 4% of its genome derived from Galla, Alpine and Toggenburg respectively. This may be attributed by noting Saanen was introduced in Kenya to crossbreed with Galla goat later than other breeds (Toggenburg and Alpine). From the admixture results, it can be concluded that Saanen goat has attained the lowest level of crossbreeding in comparison with other genotypes because the highest percentage of its genome has resulted from Galla goat. For example, according to Bett *et al* [17] Alpine has achieved up to 87.5% German Alpine blood level which is confirmed in this study the low proportion of Galla (10%) in its genome. Due to inadequate technical capacity on the farmers’ side on how to manage the rather complex crossbreeding programs, they could not be able to differentiate the change in blood level phenotypically for Saanen crossbred goats since both Saanen and Galla are white in body colour. Additionally, in recent years, it has been noted that some dairy goat breeds have spread to other areas apart from the original entry areas [15, 43]. This is in agreement with what came out from household survey that Saanen farmers have been purchasing Alpine and Toggenburg bucks/does for breeding due to lack of Saanen breeding bucks/does [44].

On the other hand, Alpine and Toggenburg goats shared some links with the Galla goat; 10% and one percent (1%), respectively. This was expected because Galla goat was used as the founder population for crossbreeding with Saanen, Alpine and Toggenburg breeds [12, 15, 45].

It is worthwhile noting that Toggenburg was the least admixed genotype in this study, this could be because of well-controlled breeding program practised by farmers under Meru Dairy Goat Breeders Association of Kenya (MDGBA). Members of the association are only allowed to use buck for breeding that belongs to the association.

## Conclusion and recommendations

The study revealed some levels of genetic diversity between goat genotypes in Kenya. Clear genetic divergence between goats in this study was identified which suggested distinct genetic resources in goat genotypes that should be sustainably utilized and conserved. The results also discovered that, there is gene-flow from exotic goats (Alpine, Toggenburg and Saanen) to the Galla populations due to crossbreeding program practised resulting in admixed goat genotypes. It is important to strengthen the various dairy goat breeders associations because urgent management efforts are essential to improve productivity, utilize and conserve the various goat genetic resources. The information generated from this study forms the basis for future genetic resource improvement programs in goat genotypes in Kenya.

## Acknowledgments

The authors wish to sincerely acknowledge Kenya Agricultural and Livestock Research Organizations (KALRO) for granting the study leave for the first author and the Egerton University (Njoro, Kenya) for providing support to undertake the study. We express our thanks to Dr. Winfred Mutisya and Dr. Ndiritu Nyaga for assisting in collecting the whole blood following FAO (2011) guidelines. We are grateful to Leonard Okutoyi and Angela M’kwenda from KALRO-Biotechnology Kabete for assisting in extraction, performing the quality and quantity measurements on the genomic DNA samples. The dairy goat farmers and Naivasha sheep and goat breeding station for availing their animals for sample collection are importantly honoured.

